# The unfolded protein response of the endoplasmic reticulum protects *Caenorhabditis elegans* against DNA damage caused by stalled replication forks

**DOI:** 10.1101/2023.02.23.529758

**Authors:** Jiaming Xu, Brendil Sabatino, Stefan Taubert

## Abstract

All animals must maintain genome and proteome integrity, especially when experiencing endogenous or exogenous stress. To cope, organisms have evolved sophisticated and conserved response systems: unfolded protein responses (UPRs) ensure proteostasis while DNA damage responses (DDRs) maintains genome integrity. Emerging evidence suggests that UPRs and DDRs crosstalk, but this remains poorly understood. Here, we demonstrate that depletion of the DNA primases *pri-1* or *pri-2*, which synthesize RNA primers at replication forks and whose inactivation causes DNA damage, activates the UPR of the endoplasmic reticulum (UPR-ER) in *Caenorhabditis elegans*, with especially strong activation in the germline. We observed activation of both the inositol-requiring-enzyme 1 (*ire-1*) and the protein kinase RNA-like ER kinase (*pek-1*) branches of the UPR-ER. Interestingly, activation of the UPR-ER output gene heat shock protein 4 (*hsp-4*) was partially independent of its canonical activators, *ire-1* and X-box binding protein (*xbp-1*), and instead required the third branch of the UPR-ER, activating transcription factor 6 (*atf-6*), suggesting functional redundancy. We further found that primase depletion specifically induces the UPR-ER, but not the distinct cytosolic or mitochondrial UPRs, suggesting that primase inactivation causes compartment-specific rather than global stress. Functionally, loss of *ire-1* or *pek-1* sensitized animals to replication stress caused by hydroxyurea. Finally, transcriptome analysis of *pri-1* embryos revealed several deregulated processes that could cause UPR-ER activation, including protein glycosylation, calcium signaling, and fatty acid desaturation. Together, our data show that the UPR-ER, but not other UPRs, responds to replication fork stress and that the UPR-ER is required to alleviate this stress.

## Introduction

The endoplasmic reticulum (ER) of eukaryotes is a dynamic membrane network required for many cellular processes, and ER homeostasis is critical to cellular and organismal health. ER homeostasis is disturbed by both impaired proteostasis and by ER membrane lipid disequilibrium (Gardner *et al*. 2013; Senft and Ronai 2015; Xu and Taubert 2021; Celik *et al*. 2023). To ensure ER function and cell viability, a conserved adaptive mechanism has evolved that restores ER homeostasis during stress: the ER unfolded protein response (UPR-ER) (Walter and Ron 2011; Gardner *et al*. 2013; Senft and Ronai 2015; Hetz *et al*. 2020; Xu and Taubert 2021). In higher eukaryotes, the UPR-ER consists of three parallel ER stress sensing and transducing branches: the Inositol-Requiring-Enzyme 1 (IRE1/IRE-1, also known as Endoplasmic Reticulum to Nucleus signaling 1 or ERN1 in mammals) branch (Adams *et al*. 2019), the protein kinase RNA-like ER kinase (PERK/PEK-1, also known as Eukaryotic Translation Initiation Factor 2α Kinase 3 or EIF2AK3) branch (McQuiston and Diehl 2017), and the Activating Transcription Factor 6 (ATF6/ATF-6) branch (Hillary and FitzGerald 2018). Together, they alleviate ER stress by reprogramming transcription and translation to promote protein folding, degradation, and transport, as well as lipid synthesis and remodeling. If ER stress cannot be resolved, the UPR-ER switches from promoting survival and adaptation to triggering apoptosis, ensuring tissue and organism integrity (Walter and Ron 2011; Hetz *et al*. 2020).

Like ER homeostasis, genome integrity is paramount for cellular and organismal health. Cells have to safeguard against DNA damage caused by endogenous and exogenous agents that induce different types of DNA lesions. To repair and mitigate DNA damage, cells activate DNA repair pathways and modulate cell cycle progression and apoptosis, a response collectively known as the DNA damage response (DDR) (Jackson and Bartek 2009; Gartner and Engebrecht 2021; McClure *et al*. 2022). Repairing this damage is critical to ensuring faithful DNA replication and thus cell division and organism development, growth, maintenance, and aging.

Interestingly, there is crosstalk between the UPR-ER and the DDR (González-Quiroz *et al*. 2020; Bolland *et al*. 2021). In yeast, *IRE1* promotes DNA repair via several different pathways, and deletion of *IRE1* sensitizes yeast to genotoxic stress and causes chromosome loss even in unstressed conditions (Henry *et al*. 2010). Moreover, activation of the checkpoint pathway by DNA damage upregulates the key UPR-ER output transcription factor Hac1p (Tao *et al*. 2011). Similarly, in cultured human cell lines, IRE-1 promotes genome integrity through the downstream effector X-box binding protein 1 (XBP1; the human ortholog of Hac1p), which directly regulates several DDR pathways, and through non-canonical regulated IRE-1-dependent decay (RIDD) of mRNA (Dufey *et al*. 2020). Consequently, loss of XBP1 correlates with increased DNA damage (Acosta-Alvear *et al*. 2007; Argemí *et al*. 2017; Lyu *et al*. 2019; González-Quiroz *et al*. 2020). Moreover, DNA damaging agents such as camptothecin and ionizing radiation trigger UPR-ER activation in cancer cell lines through the conserved DDR sensor ATM (Hotokezaka *et al*. 2020), and the genome integrity regulator p53 mediates ER structure remodeling in chemically induced genotoxicity (Zheng *et al*. 2018). The UPR-ER, especially the IRE-1 branch, is therefore both activated by DNA damage and functionally required to repair such damage.

Whether integration of the DDR and the UPR-ER also occurs in animals, and how different tissues respond in this context is less well understood. In the nematode worm *Caenorhabditis elegans*, DNA damage in the adult germline promotes stress resistance in the postmitotic soma via MAP kinase signaling, innate immune responses, and the ubiquitin proteasome system (Ermolaeva *et al*. 2013), which, like the UPR-ER, maintains proteostasis (Papaevgeniou and Chondrogianni 2014; Zhang *et al*. 2022). Moreover, *C. elegans xbp-1* is required to express DNA repair genes (Shen *et al*. 2005). Recently, increased stress resistance to the ER stressors dithiothreitol (DTT) and tunicamycin was observed in *C. elegans* exposed to UV, which increased the activity of the IRE-1–XBP-1 branch by elevating the levels of unsaturated phosphatidylcholine (Deng *et al*. 2021), a key ER membrane lipid (Meer *et al*. 2008). However, this study predominantly analyzed DDR and UPR-ER signaling in the *glp-1* mutant, which lacks a germline, is long-lived and stress resistant, and shows dysregulation of DDR genes (Arantes-Oliveira 2002; Boyd *et al*. 2010; TeKippe and Aballay 2010; Ratnappan *et al*. 2014; Goh *et al*. 2018). The interaction of the UPR-ER and the DDR in wild-type animals with an intact germline, which is the primary tissue of active DNA repair, therefore remains incompletely understood.

In a screen for genes whose inactivation causes UPR-ER activation in wild-type *C. elegans*, we identified two DNA primase genes (Ho *et al*. 2020). Eukaryotic primase complexes synthesize short RNA primers required for initiating lagging strand DNA replication and also contribute to DNA repair and possibly transcription (Guilliam *et al*. 2015; Yoon *et al*. 2018). Abnormal primase function causes stalled replication forks, leading to DNA damage and genome instability. This therefore provided us with the opportunity to study the relationship between primase function, replication stress, and the UPR-ER in animals with an intact germline. Here, we show that primase inactivation and UV-C irradiation activate both the IRE-1 and the PEK-1 branches of the UPR-ER, with stronger induction in the germline than in the soma. Interestingly, activation of the *hsp-4* gene, which canonically requires the IRE-1–XBP-1 axis, required *atf-6* in the germline, suggestion differential regulatory mechanisms. We further found that primase inactivation selectively activated the UPR-ER but not the cytosolic or mitochondrial UPRs, arguing for a specific role of the UPR-ER in maintaining genome integrity. We also showed that loss of *ire-1* or *pek-1* sensitizes *C. elegans* to replication stress, showing that the UPR-ER is functionally protective. RNA-seq analysis revealed several pathways comprising both proteostasis and lipidostasis that could underlie UPR-ER activation following replication stress. Collectively, our data show that the UPR-ER plays important roles in ensuring genome integrity in *C. elegans*.

## Materials and Methods

### Worm strains

The following worm strains were used: N2 wild-type, SJ4005 *zcIs4 [hsp-4p::gfp] V* (Calfon *et al*. 2002), SJ17 *xbp-1(zc12) III;zcIs4 [hsp-4p::gfp] V* (Calfon *et al*. 2002), SJ30 *ire-1(zc14) II;zcIs4 [hsp-4p::gfp] V* (Calfon *et al*. 2002), SJ4100 *zcIs13 [hsp-6p::gfp] V* (Yoneda *et al*. 2004), TJ375 *gpIs1 [hsp-16*.*2p::gfp]* (Henderson and Johnson 2001), *Patf-4(uORF)::gfp::unc-54(3’UTR)* (Venz *et al*. 2020), *xbp-1(tm2482) III* (Richardson *et al*. 2011), RB545 *pek-1(ok275) V* (Consortium 2012), RB925 *ire-1(ok799) II* (Consortium 2012), RB772 *atf-6(ok551) X* (Consortium 2012), PHX2824 *hsp-4::gfp(syb2824) II* (generated by SunyBiotech Co., Ltd., Fujian, China), and STE142 *hsp-4::gfp(syb2824) II*; *atf-6(ok551) X* (this study). All strains were backcrossed six times to the laboratory N2 wild-type background prior to use.

### Worm growth conditions

We cultured *C. elegans* strains at 20°C on nematode growth medium (NGM)-lite agar plates with *E. coli* OP50 as food source, except for RNA interference (RNAi), for which HT115 strain was used (Ho *et al*. 2020). To developmentally synchronize worm populations, gravid adult worms were treated with alkaline sodium hypochlorite solution to extract embryos, which were washed twice with M9, and then plated onto an unseeded NGM-lite plate to allow hatching overnight. When imaging worms, adult worms were bleached >10 minutes until auto-fluorescent mother bodies disappeared. The resulting synchronized L1 larvae were transferred onto OP50 NGM-lite plates or RNAi plates (NGM-lite plates containing 25 μg/mL carbenicillin (BioBasic CDJ469), 2 mM IPTG (Santa Cruz sc-202185B), and 12.5 μg/mL tetracycline (BioBasic TB0504). RNAi plates were seeded twice with the appropriate HT115 RNAi bacteria (Ahringer library, Source BioScience); RNAi clones were sequenced prior to use to ensure construct identity. Synchronized L1 worms were placed on RNAi plates and grown until they reached the desired developmental stage.

### DIC and fluorescence microscopy

Worms were mounted onto 2% (w/v) agarose pads containing a drop of 20 mM sodium azide (NaN_3_) for microscopy. Eggs were picked from plates onto 2% (w/v) agarose pads containing a drop of M9 for microscopy. Worms were imaged using differential interference contrast (DIC) and fluorescence optics through a CoolSnap HQ camera (Photometrics, Tucson, AZ, USA) on a Zeiss Axioplan 2 compound microscope (Carl Zeiss Microscopy, Thornwood, NY, USA). All GFP images were taken at the same exposure time (300ms). Using the ImageJ software, the images in the GFP channel were adjusted to the same brightness (maximum display value = 4095, minimum = 201; these parameters were applied to all GFP images used for quantification in this study) and contrast levels for subsequent display and quantification purposes. Analysis of overall fluorescence intensity of individual worms was performed by tracing the outline of worms on the corresponding DIC images, and then normalized for area and background fluorescence, as described (Shomer *et al*. 2019).

### Protein extraction and immunoblots

Whole-worm protein extracts were generated by sonication in radioimmunoprecipitation assay (RIPA) lysis buffer with cOmpleteTM Protease Inhibitor Cocktail (Roche #4693116001) and β-glycerophosphate (Sigma-Aldrich G6251). Protein concentrations were determined using the RC DC Protein Assay kit (Bio-Rad #500-0121), and sodium dodecyl-sulfate polyacrylamide gel electrophoresis (SDS-PAGE) analysis and immunoblotting were performed as described (Hou *et al*. 2014), using anti Ser51-Phospho-eIF2α rabbit antibody (Cell Signaling Technologies #9721), anti-α-tubulin mouse antibody (Sigma #T9026), and anti-rabbit HRP conjugated (NEB #7074) and anti-mouse HRP conjugated (Cell Signaling Technologies #7076) secondary antibodies. Detection was done using ECL (Pierce #32109).

### Exposure to genotoxic agents

For UV-C exposure, synchronized populations of day-1 adult *C. elegans* were placed on NGM-lite plates seeded with a thin layer of OP50. Uncovered NGM-lite plates were then placed in a Stratalinker 2400 UV Crosslinker (Stratagene) and irradiated with wavelength 254 nm light at 400J/m^2^. After 24hr of recovery at 20°C, worms and embryos were mounted and imaged.

For hydroxyurea (HU) exposure L1 recovery experiments, age-synchronized L1 populations were grown for 72hr on NGM-lite or on NGM-lite containing either 5 or 10mM HU (Sigma-Aldrich H8627). Then, worms were transferred to OP50-seeded NGM-lite plates for recovery and egg laying. After 4hr, the number of eggs was counted for each genotype and condition, as indicated.

For HU exposure L4 recovery experiments, synchronized L1 worms were grown for 48hr. Then, age-synchronized L4 populations were transferred to and maintained on either NGM-lite plates or NGM-lite plates containing 20mM HU for 24hr. Then, adult worms were transferred to OP50 plates for recovery and egg laying. After 4hr, the number of eggs for each genotype and condition was counted.

To measure developmental rate, synchronized L1 populations were grown for 48hr on NGM-lite plates containing DMSO vehicle or 15mM HU. Then, the number of L4 or older worms and the total number of worms were counted for each genotype and condition.

For body size quantification, synchronized L1 stage worms were grown for 72hr on NGM-lite plates containing DMSO or 15mM HU, before >10 worms for each genotype and condition were imaged.

### RNA-sequencing and data analysis

Synchronized L1 N2 worm populations fed empty vector (EV) or *pri-1* RNAi were grown for 96 hours at 20°C and allowed to lay eggs. Plates were washed with M9 twice to remove adults and hatched worms before eggs were harvested with a cell scraper. The collected eggs were washed twice with M9 to remove bacteria, and then flash-frozen in an ethanol-dry ice bath. For total RNA extraction, eggs were thawed in Trizol and sonicated. Total RNA was extracted using Trizol and 1-bromo-3-chloropropane, as described (Doering *et al*. 2022). RNA integrity and quantity were assessed on an Agilent Technologies 2100 Bioanalyzer System.

Library preparation and sequencing was performed by The Center for Applied Genomics, SickKids, Toronto, ON (http://www.tcag.ca). Briefly, RNA was prepared for sequencing using the NEBNext® Ultra II Directional RNA Library Prep Kit for Illumina (NEB #E7760). Sequencing was performed on an Ilumina NovaSeq 6000 instrument equipped with an S4 flow cell generating 150bp paired-end reads. Low quality reads and adapter sequences were trimmed using trimmomatic 0.36 (Bolger *et al*. 2014) with parameters LEADING:3 TRAILING:3 SLIDINGWINDOW:4:15 MINLEN:36. Trimmed reads were quantified to the *C. elegans* Ensembl transcriptome build WBcel235 using Salmon ver1.4.0 (Patro *et al*. 2017) with parameters -l A -p 8 --gcBias --validateMappings. Then, transcript-level read counts were imported into R and summed into gene-level read counts using tximport (Soneson *et al*. 2016) (genes listed in Supplementary Table S1). Genes not expressed at a level greater than 10 reads in at least three of the samples were excluded from further analysis. Differential expression analysis was performed with quasi-likelihood F-test with the generalized linear model (GLM) approach in edgeR (Robinson *et al*. 2010). Genes with pvalue <0.05 and FDR<0.05 were considered differentially expressed. RNA-seq data were deposited in Gene Expression Omnibus under the accession number GSE225569. Gene Set Enrichment Analysis (GSEA) using either the Biological Process (BP (Chagoyen and Pazos 2010)) or the Kyoto Encyclopedia of Genes and Genomes (KEGG (Kanehisa 1997; Kanehisa *et al*. 2020)) as underlying databases was performed with eVITTA (Cheng *et al*. 2021), using pval< 0.05 and padj < 0.25 as cutoffs.

### Statistical analysis

p values were calculated using two-tailed Student’s t tests, Welch’s t tests, one-way ANOVA tests, or two-way ANOVA tests using GraphPad Prism 9, as reported in the figure legends. Scatter plots were generated in GraphPad Prism 9. Error bars denote standard deviation; the number of independent experiments performed and number of animals studied are indicated in the figure legends.

## Results

### Knockdown of the *C. elegans* primase genes *pri-1* or *pri-2* activates the *ire-1* branch of the UPR-ER in embryos

We previously showed that RNAi against the two DNA primase subunit genes of *C. elegans, pri-1* or *pri-2*, caused activation of the UPR-ER (Ho *et al*. 2020). To validate this finding, we quantified the induction of *hsp-4p::gfp*, a widely-used transcriptional reporter for the ER stress-inducible, *ire-1*- and *xbp-1*-activated *hsp-4* gene promoter (Calfon *et al*. 2002; Hou *et al*. 2014; Ho *et al*. 2020). We found that *pri-1* or *pri-2* RNAi induced *hsp-4p::gfp* fluorescence in the worm soma approximately 1.5-2 fold (Fig. 1A,B). Interestingly, we observed a larger increase in *hsp-4p::gfp* activity (approximately ∼4 fold) in F1 embryos from RNAi-fed P0 adults (Fig. 1C,D) This phenotype manifests despite the fact that F1 eggs from *pri-1* or *pri-2* RNAi treated P0 adults never hatch, but rather arrest at the early embryogenesis/pre-morphogenetic stage due to persisting replication fork stalling, which causes double stranded DNA breaks (DSBs) (Zeman and Cimprich 2014). UPR-ER activation in the F1 generation by *pri-1* or *pri-2* RNAi in P0 suggested a link between the UPR-ER and genotoxic stress in embryos.

**Figure 1.**
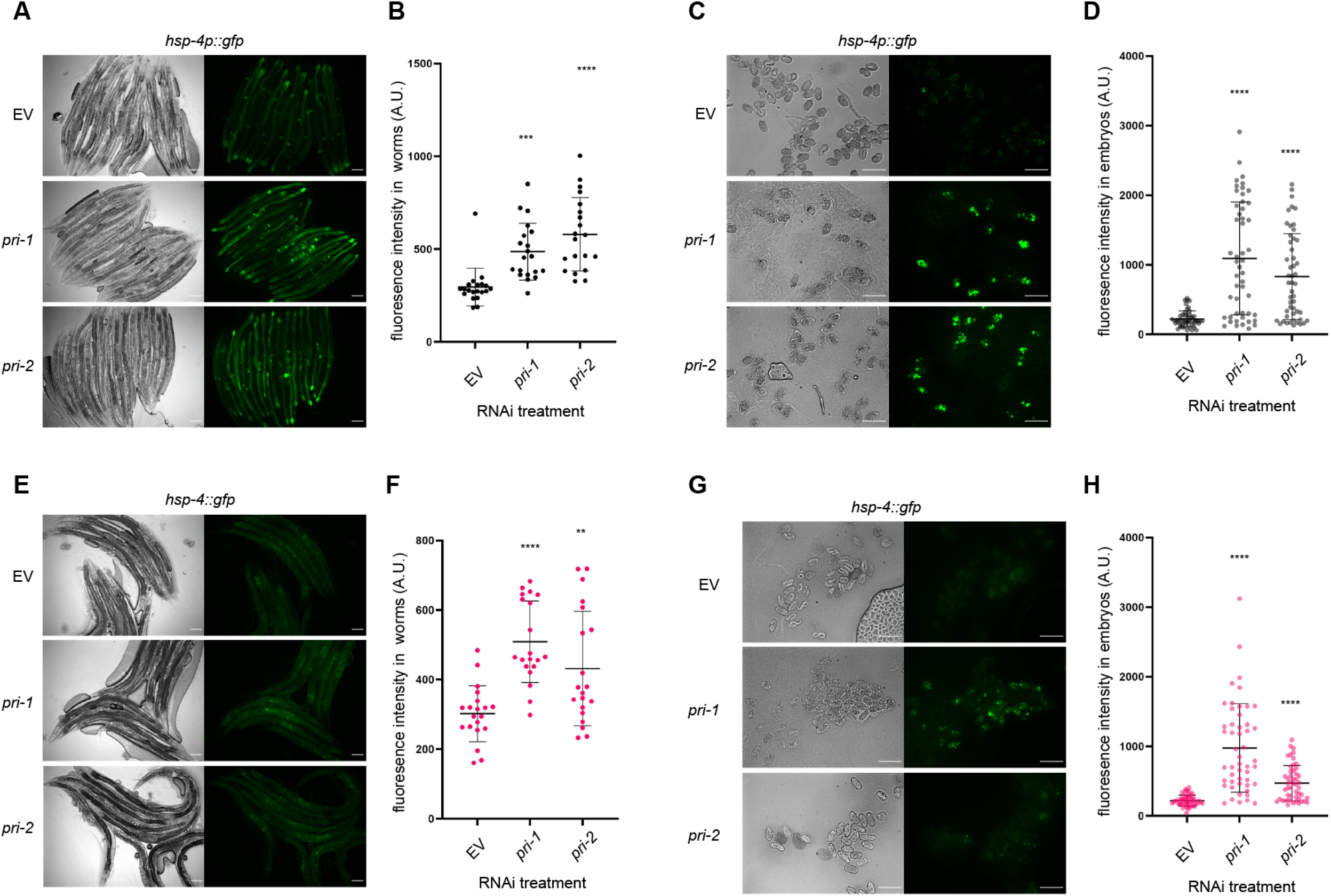
*pri-1* or *pri-2* knockdown induces IRE-1 branch activity in the soma and embryos of *C. elegans*. **(A, B)** The figure shows representative micrographs (A) and whole-worm GFP quantification (B) of *hsp-4p::gfp* adult worms fed EV, *pri-1*, or *pri-2* RNAi (n=3 experiments totaling >20 animals per RNAi treatment). **(C, D)** The figure shows representative micrographs (C) and GFP quantification (D) of F1 embryos laid by *hsp-4p::gfp* adult worms fed EV, *pri-1*, or *pri-2* RNAi (n=3 experiments totaling >50 embryos per RNAi treatment). **(E, F)** The figure shows representative micrographs (E) and whole-worm GFP quantification (F) of *hsp-4::gfp* adult worms fed EV, *pri-1*, or *pri-2* RNAi (n=2 experiments totaling >20 animals per RNAi treatment). **(G, H)** The figure shows representative micrographs (G) and GFP quantification (H) of embryos laid by *hsp-4::gfp* adult worms fed EV, *pri-1*, or *pri-2* RNAi (n=2 experiments totaling >50 embryos per treatment). In all micrographs, the scale bar represents 100µm. In dot plots, each dot represents the signal detected in one individual worm or embryo; error bars represent standard deviation. Statistical analysis for B, D, F, H: **p<0.01, ***p<0.001, ****p<0.0001 vs. EV RNAi-treated control (Brown-Forsythe and Welch ANOVA test corrected for multiple comparisons using the Dunnett T3 method).

To confirm activation of the endogenous UPR-ER by *pri-1* or *pri-2* RNAi, we studied a genome-edited strain wherein the 3’ end of the *hsp-4* coding sequence is tagged with *gfp*, resulting in a C-terminal HSP-4:: GFP fusion protein (hereafter referred to as *hsp-4::gfp*). To validate this strain, we examined GFP intensity following proteotoxic stress by tunicamycin and lipotoxic stress by *mdt-15* RNAi, both established UPR-ER inducers (Calfon *et al*. 2002; Hou *et al*. 2014). As expected, we observed elevated GFP intensity in *hsp-4::gfp* worms challenged with tunicamycin or *mdt-15* RNAi (Supplementary Fig. 1), suggesting that this strain faithfully reports on the regulation of endogenous *hsp-4* by different stresses. Next, we treated the *hsp-4::gfp* reporter strain with *pri-1* or *pri-2* RNAi and studied GFP fluorescence in the soma of P0 adult worms and in F1 embryos. We observed strong induction of endogenous HSP-4::GFP in F1 embryos, and weaker induction in P0 somatic cells (Fig. 1E-H). These data suggest that loss of primase function and subsequent replication defects trigger UPR-ER activation in embryos and in somatic cells of *C. elegans*, with stronger induction in embryos.

### Knockdown of *pri-1* or *pri-2* induces embryonic *hsp-4* partially independently of *ire-1* and *xbp-1*

Canonical *hsp-4* induction requires the transmembrane ER stress sensor *ire-1* and the downstream transcription factor *xbp-1* (Shen *et al*. 2001; Richardson *et al*. 2010). Thus, we studied *hsp-4* induction in *ire-1;hsp-4p::gfp* and *xbp-1;hsp-4p::gfp* worms treated with *pri-1* or *pri-2* RNAi. Consistent with canonical UPR-ER induction in somatic cells, increased fluorescence in *pri-1* or *pri-2* treated worms depended completely on *ire-1* and *xbp-1* (Fig. 2A,B). In contrast, significant induction of *hsp-4p::gfp* remained in embryos when *ire-1* or *xbp-1* was deleted (Fig. 2C,D). This suggests that additional genes are required to induce *hsp-4* in embryos experiencing replication stress.

**Figure 2.**
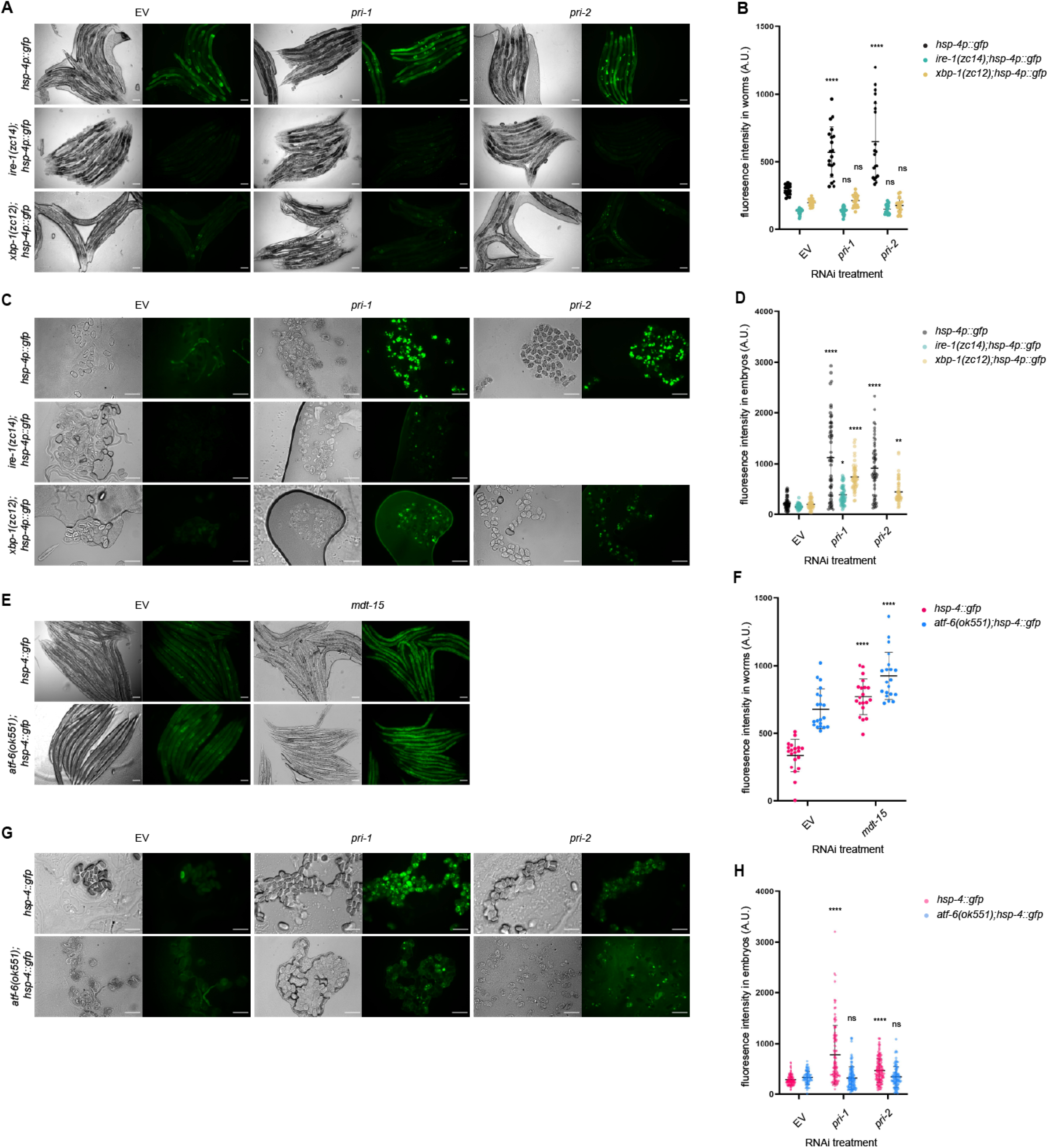
Activation of *hsp-4* by *pri-1* or *pri-2* RNAi requires *ire-1*, x*bp-1*, and *atf-6*. **(A, B)** The figure shows representative micrographs (A) and whole-worm GFP quantification (B) of *hsp-4p::gfp, ire-1(zc14);hsp-4p::gfp*, and *xbp-1(zc12);hsp-4p::gfp* adult worms fed EV, *pri-1*, or *pri-2* RNAi (n=3 experiments totaling >20 individual animals per RNAi treatment). **(C, D)** The figure shows representative micrographs (C) and GFP quantification (D) of embryos laid by *hsp-4p::gfp, ire-1(zc14);hsp-4p::gfp*, and *xbp-1(zc12);hsp-4p::gfp* adult worms fed EV, *pri-1*, or *pri-2* RNAi (n=3 experiments totaling >50 individual embryos per RNAi treatment). **(E, F)** The figure shows representative micrographs (E) and whole-worm GFP quantification (F) of *hsp-4::gfp* and *atf-6 (ok551);hsp-4::gfp* adult worms fed EV or *mdt-15* RNAi (n=3 experiments totaling >20 individual animals per RNAi treatment). **(G, H)** The figure shows representative micrographs (G) and GFP quantification (H) of embryos laid by *hsp-4::gfp* and *atf-6 (ok551);hsp-4::gfp* adult worms fed EV, *pri-1*, or *pri-2* RNAi (n=3 experiments totaling >75 individual embryos per treatment). In all micrographs, the scale bar represents 100µm. In dot plots, each dot represents the signal detected in one individual worm or embryo; error bars represent standard deviation. Statistical analysis: B, D: ns p>0.05, *p<0.05, **p<0.01, ****p<0.0001, vs. EV RNAi-treated control of the same genotype (two-way ANOVA test corrected for multiple comparisons using Sidak’s method).

### *atf-6* is required for *hsp-4* induction in *pri-1* or *pri-2* RNAi treated embryos

In mammals, ATF6 is required for the transcription of *XBP1* mRNA (Yoshida *et al*. 2001; Lee *et al*. 2002). Thus, we tested if *C. elegans atf-6* is required for replication stress-induced *hsp-4* induction. We crossed the *hsp-4::gfp* translational reporter into a strain bearing the *atf-6(ok551)* null allele, and treated *atf-6;hsp-4::gfp* worms with *pri-1* or *pri-2* RNAi. As a control, we first studied *mdt-15* RNAi, which caused *hsp-4::gfp* induction in the soma despite the *atf-6* mutation (Fig. 2E,F), suggesting that *hsp-4* induction in somatic tissues does not require *atf-6*; unexpectedly, *atf-6* loss alone induced *hsp-4* expression in the soma (Fig. 2E,F). Notably, *atf-*6 deletion reduced the increased fluorescence in embryos treated with *pri-1* or *pri-2* RNAi (Fig. 2G,H). This indicates that *hsp-4* induction in *pri-1* or *pri-2* RNAi treated embryos depends on *atf-6*.

### *pri-1* or *pri-2* knockdown activates the *pek-1* branch of the UPR-ER

The animal UPR-ER also features the *pek-1* branch (McQuiston and Diehl 2017), so we tested if *pri-1* or *pri-2* RNAi activate *Patf-4(uORF)::gfp*, a reporter of *pek-1* branch activity (Venz *et al*. 2020). We found that *pri-1* or *pri-2* RNAi robustly induced *Patf-4(uORF)::gfp* in F1 embryos (Fig. 3A,B), with weaker, but significant induction in the soma of P0 worms (Fig. 3C,D). As an additional readout, we performed Western blots on embryos to detect phospho-Ser51 on eIF2α, another marker for activated PEK-1 (Nukazuka *et al*. 2008). We observed increased levels of phospho-Ser51 in *pri-1* or *pri-2* RNAi treated embryos (Fig. 3E, Supplementary Fig. 2). Thus, *pri-1* or *pri-2* RNAi activates both the *ire-1* and *pek-1* branches of the UPR-ER in the soma and in embryos.

**Figure 3.**
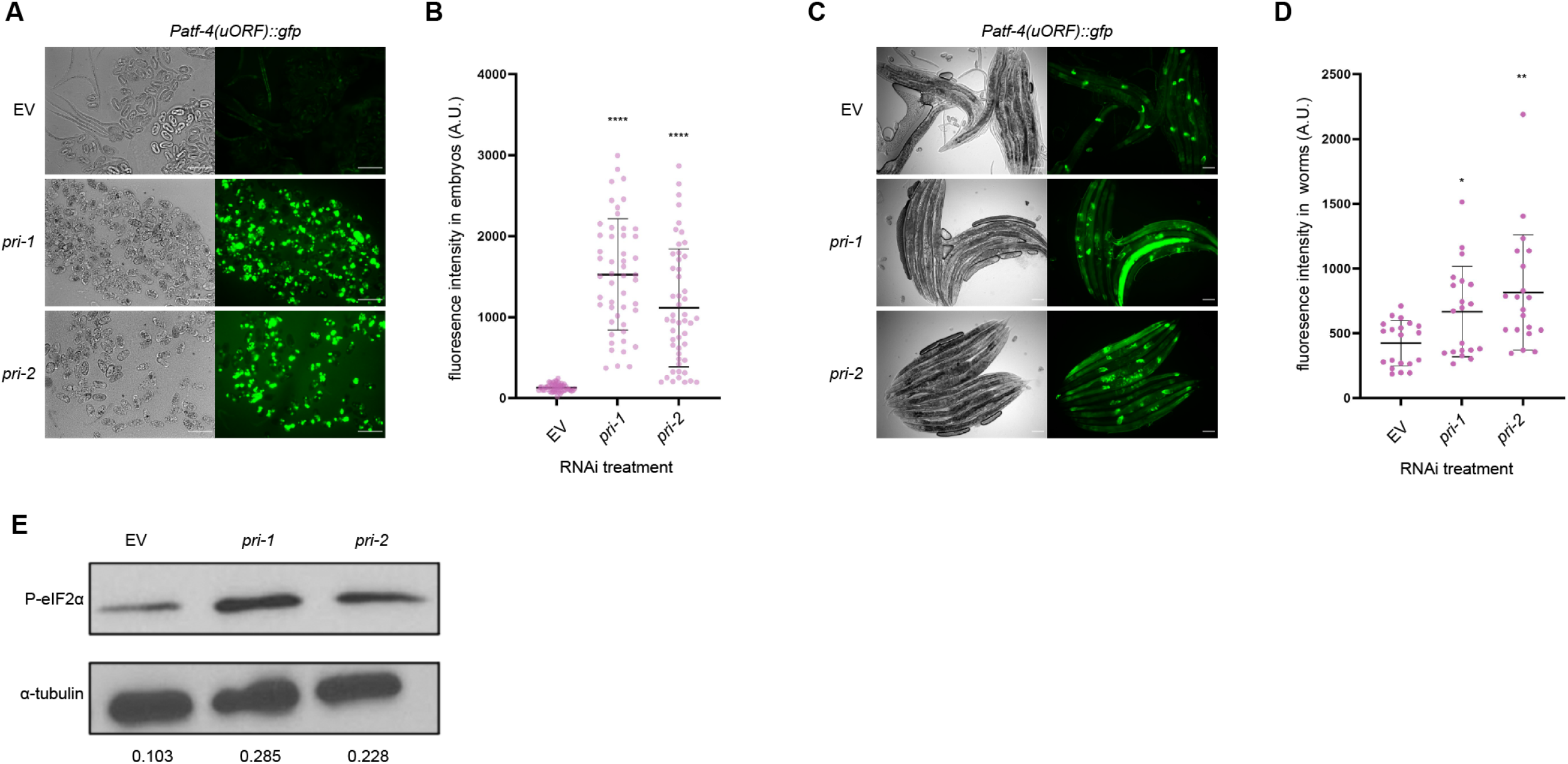
*pri-1* or *pri-2* RNAi activate the PEK-1 branch in the soma and embryos of *C. elegans*. **(A, B)** The figure shows representative micrographs (A) and GFP quantification (B) of embryos laid by *Patf-4(uORF)::gfp* adult worms fed EV, *pri-1*, or *pri-2* RNAi (n=3 experiments totaling >50 individual embryos per RNAi treatment). **(C, D)** The figure shows representative micrographs (C) and whole-worm GFP quantification (D) of *Patf-4(uORF)::gfp* adult worms fed EV, *pri-1*, or *pri-2* RNAi (n=3 experiments totaling >20 individual animals per RNAi treatment). In all micrographs, the scale bar represents 100µm. In dot plots, each dot represents the signal detected in one individual worm or embryo; error bars represent standard deviation. Statistical analysis for B, D: *p<0.05, **p<0.01, ****p<0.0001 vs. EV RNAi-treated control (Brown-Forsythe and Welch ANOVA test corrected for multiple comparisons using the Dunnett T3 method). **(E)** The immunoblot depicts the levels of phospho-Ser51 eIF2α (P-eIF2α) and α-tubulin in EV, *pri-1*, or *pri-2* RNAi treated worm embryos (n=2-3, for additional repeats please see Supplementary Figure S2). Numbers represent intensity of the P-eIF2α bands normalized to the corresponding α-tubulin bands.

### The cytosolic and the mitochondrial UPRs are not substantially induced by *pri-1* or *pri-2* knockdown

Induction of the UPR-ER due to *pri-1* or *pri-2* RNAi might reflect general protein misfolding in terminally arrested embryos. Thus, we monitored the activity of the cytosolic and mitochondrial UPRs with their well-established reporters, *hsp-16*.*2p::gfp* and *hsp-6p::gfp*, respectively (Rea *et al*. 2005; Bennett *et al*. 2014). In the worm soma, heat shock strongly induced *hsp-16*.*2p::gfp* (Supplementary Fig. 3A,B), and positive control *cco-1* RNAi strongly induced *hsp-6p::gfp* (Fig. 4A,B), as expected (Bennett *et al*. 2014). In contrast, although *pri-1* or *pri-2* RNAi effectively induced *hsp-4p::gfp* (Fig. 4C,D), they did not induce *hsp-6p::gfp* or *hsp-16*.*2p::gfp* in somatic tissues; activity of both reporters was in fact reduced (Fig. 4A,B,E,F). Similarly, whereas heat stress strongly (approximately five-fold) induced *hsp-16*.*2p::gfp* throughout the embryo (Supplementary Fig. 3C,D), *pri-1* or *pri-2* RNAi only weakly (less than two-fold), albeit still significantly, induced *hsp-16*.*2p::gfp* fluorescence in embryos (Fig. 4G,H), while strongly inducing *hsp-4p::gfp* (Fig. 4I,J). Thus, *pri-1* or *pri-2* RNAi-induced replication stress appears to predominantly trigger the UPR-ER.

**Figure 4.**
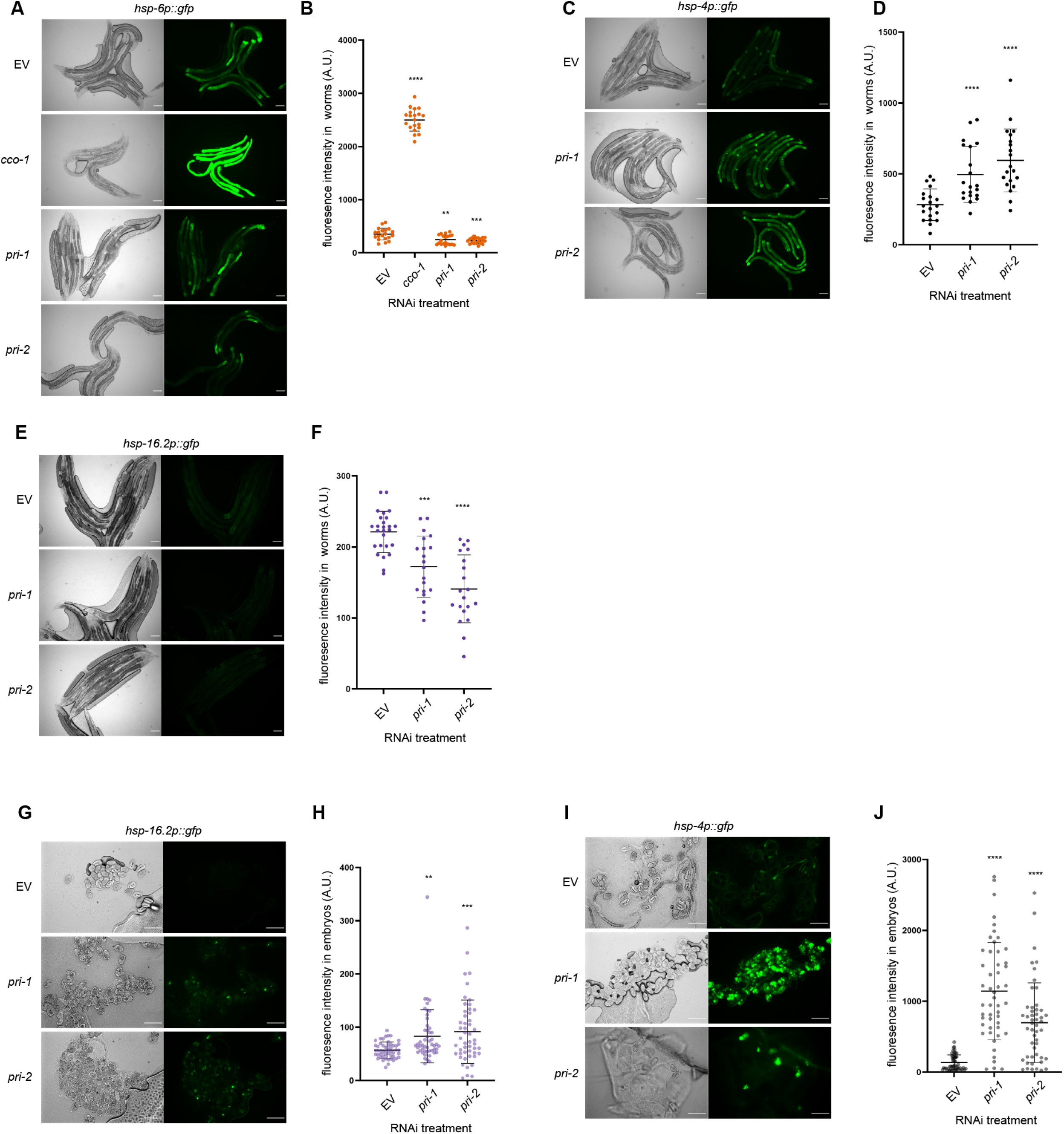
*pri-1* or *pri-2* RNAi preferentially induces the UPR-ER. **(A, B)** The figure shows representative micrographs (A) and whole-worm GFP quantification (B) of *hsp-6p::gfp* adult worms fed EV, *cco-1, pri-1*, or *pri-2* RNAi (n=3 experiments totaling >20 individual animals per RNAi treatment). **(C, D)** The figure shows representative micrographs (A) and whole-worm GFP quantification (B) of *hsp-4p::gfp* adult worms fed EV, *pri-1*, or *pri-2* RNAi (n=3 experiments totaling >20 individual animals per RNAi treatment). **(E, F)** The figure shows representative micrographs (E) and whole-worm GFP quantification (F) of *hsp-16*.*2p::gfp* adult worms fed EV, *pri-1*, or *pri-2* RNAi (n=2 experiments totaling >20 individual animals per RNAi treatment). **(G, H)** The figure shows representative micrographs (G) and GFP quantification (H) of embryos laid by *hsp-16*.*2p::gfp* adult worms fed EV, *pri-1*, or *pri-2* RNAi (n=3 experiments totaling >50 individual embryos per treatment). **(I, J)** The figure shows representative micrographs (I) and GFP quantification (J) of embryos laid by *hsp-4p::gfp* adult worms fed EV, *pri-1*, or *pri-2* RNAi (n=3 experiments totaling >50 individual embryos per treatment). In all micrographs, the scale bar represents 100µm. In dot plots, each dot represents the signal detected in one individual worm; error bars represent standard deviation. Statistical analysis for B, D, F, H, J: **p<0.01, ***p<0.001, ****p<0.0001 vs. EV RNAi-treated control (Brown-Forsythe and Welch ANOVA test corrected for multiple comparisons using the Dunnett T3 method).

### Inactivation of other polymerase α primase complex genes phenocopies *pri-1* or *pri-2* RNAi

Four genes encode *C. elegans* primase complex subunits: the DNA polymerase α catalytic subunit gene *pola-1*, the DNA polymerase α accessory subunit gene *div-1*, and the primase subunit genes *pri-1* and *pri-2* (Guilliam *et al*. 2015; Yoon *et al*. 2018). We asked if RNAi knockdown of *pola-1* or *div-1* phenocopied *pri-1* or *pri-2* RNAi. Indeed, *pola-1* and *div-1* RNAi activated *hsp-4p::gfp* in both the soma of P0 worms and in F1 embryos, with stronger activation in embryos than in the soma (Fig. 5A-D). This suggests that UPR-ER induction likely results from replication stress caused by defective polymerase α primase complex function.

**Figure 5.**
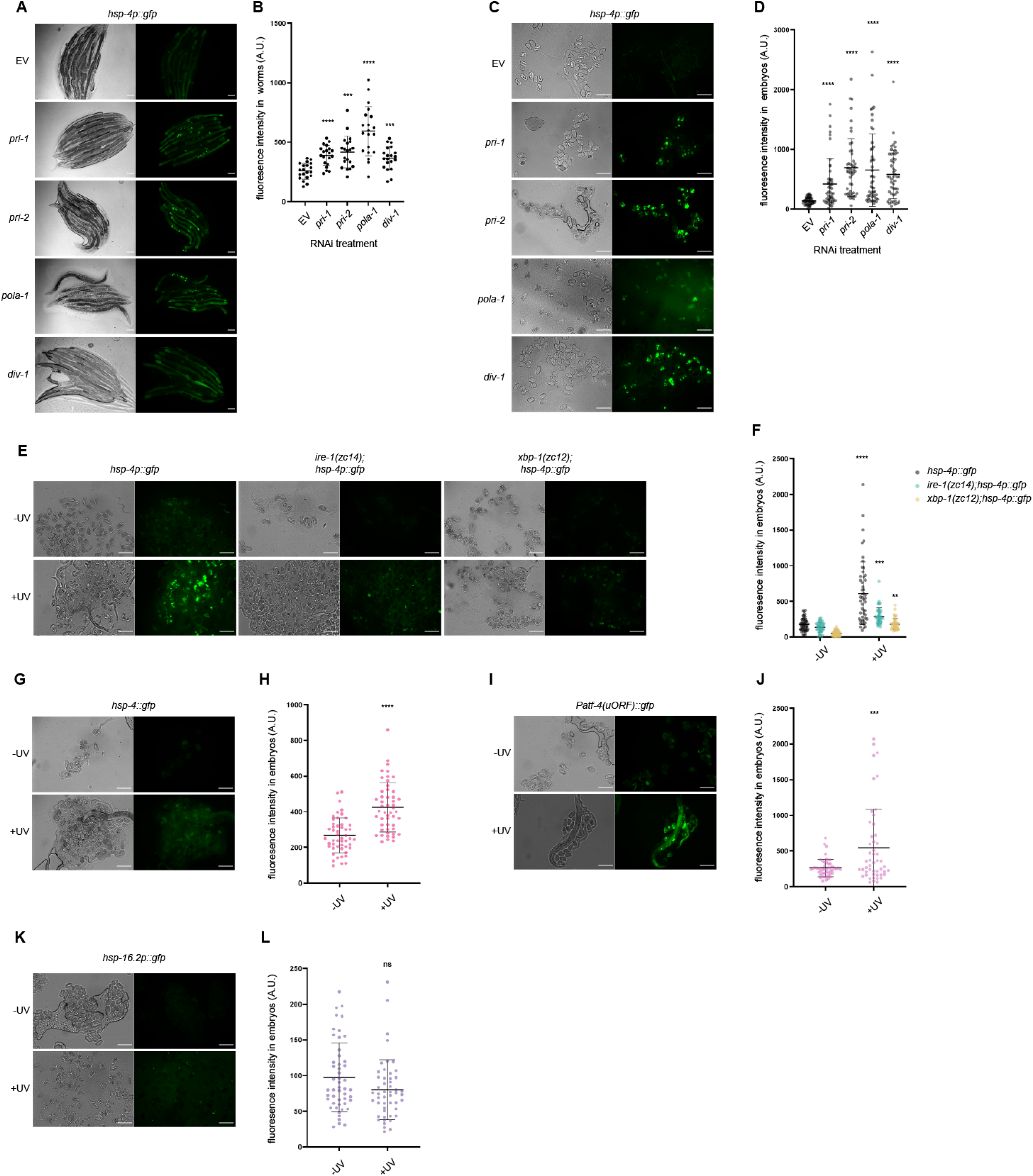
Knockdown of polymerase α primase complex subunits and UV-C irradiation cause UPR-ER activation. **(A, B)** The figure shows representative micrographs (A) and whole-worm GFP quantification (B) of *hsp-4p::gfp* adult worms fed EV, *pri-1, pri-2, pola-1*, or *div-1* RNAi (n=3 experiments totaling >20 individual animals per RNAi treatment). **(C, D)** The figure shows representative micrographs (C) and GFP quantification (D) of embryos laid by *hsp-4p::gfp* adult worms fed EV, *pri-1, pri-2, pola-1*, or *div-1* RNAi (n=3 experiments totaling >50 individual embryos per RNAi treatment). **(E, F)** The figure shows representative micrographs (E) and GFP quantification (F) of embryos laid by *hsp-4p::gfp, ire-1(zc14);hsp-4p::gfp*, and *xbp-1(zc12);hsp-4p::gfp* adult worms irradiated with 400J/m^2^ UV-C (n=3 experiments repeats totaling >50 individual embryos for each sample). **(G, H)** The figure shows representative micrographs (G) and GFP quantification (H) of embryos laid by *hsp-4::gfp* adult worms irradiated with 400J/m^2^ UV-C (n=3 experiments totaling >50 individual embryos for UV-irradiated and non-irradiated samples). **(I, J)** The figure shows representative micrographs (I) and GFP quantification of embryos laid by *Patf-4(uORF)::gfp* adult worms irradiated with 400J/m^2^ UV-C (n=3 experiments totaling >50 individual embryos for UV-irradiated and non-irradiated samples). **(K, L)** The figure shows representative micrographs (K) and GFP quantification (L) of embryos laid by *hsp-16*.*2p::gfp* adult worms irradiated with 400J/m^2^ UV-C (n=3 experiments totaling >50 individual embryos for UV-irradiated and non-irradiated samples). In all micrographs, the scale bar represents 100µm. In dot plots, each dot represents the signal detected in one individual worm or embryo; error bars represent standard deviation. Statistical analysis: B, D: ***p<0.001, ****p<0.0001 vs. EV RNAi control (Brown-Forsythe and Welch ANOVA test corrected for multiple comparisons using the Dunnett T3 method); F: **p<0.01, ***p<0.001, ****p<0.0001 vs. non-irradiated embryos of the same genotype (ordinary two-way ANOVA test corrected for multiple comparisons using Sidak’s method); H, J, L: ns p>0.05, ***p<0.001, ****p<0.0001 vs. non-irradiated embryos (Welch’s t test).

### UV-C treatment phenocopies *pri-1* or *pri-2* RNAi

Like replication block, ultraviolet C (UV-C) light induces DSBs if the resulting bipyrimidine photoproducts are not resolved by the nucleotide excision repair (NER) pathway (Stergiou *et al*. 2011). Thus, we tested if UV-C exposure in early embryos phenocopies *pri-1* or *pri-2* RNAi treatment. We irradiated day 1 P0 adult worms with 400J/m^2^ UV-C and studied F1 embryos 24 hrs thereafter. Like *pri-1* or *pri-2* RNAi, UV-C treatment strongly activated *hsp-4p::gfp* (Fig. 5E,F), in line with previously published observations (Deng *et al*. 2021). As observed for *pri-1* or *pri-2* RNAi, UV-C-induced *hsp-4p::gfp* upregulation was partially independent of *ire-1* and *xbp-1* in embryos (Fig. 5E,F). The *hsp-4::gfp* translational reporter was also induced by UV-C (Fig. 5G,H). We also observed induction of the *Patf-4(uORF)::gfp* reporter, indicating activation of the *pek-1* branch (Fig. 5I,J). In contrast, the cytosolic UPR reporter *hsp-16*.*2p::gfp* was not activated (Fig. 5K,L), suggesting that UV-C specifically activates the UPR-ER in embryos. In the soma, UV-C caused activation of the *pek-1* branch reporter *Patf-4(uORF)::gfp*; we also observed approximately two-fold activation of *hsp-16*.*2p::gfp*, whereas *hsp-4p::gfp, hsp-4::gfp*, and *hsp-6p::gfp* were not activated (Supplementary Fig. 4). Collectively, these observations suggest that UV-C irradiation, like *pri-1* or *pri-2* RNAi, cause DSBs and UPR-ER activation, especially in the embryo.

### Inactivating components of the DNA repair machinery does not activate the UPR-ER in somatic cells

The above data raised the possibility that genotoxic stress in general activates the UPR-ER. To test this hypothesis, we used RNAi to inactivate several DNA repair genes, which should cause increased DNA damage, specifically: *msh-2* (mismatch repair (Degtyareva *et al*. 2002)), *xpf-1* (nucleotide excision repair (Saito *et al*. 2009)), *him-1* (a cohesin, whose loss results in chromosomal segregation defects in mitosis and meiosis (Chan *et al*. 2003)), *mus-81* (replicative repair (O’Neil *et al*. 2013)), *dog-1* and *him-6* (whose loss causes formation of R-loops or G4 structures, causing deletions in poly-G tracts and genome instability (Cheung *et al*. 2002; Youds *et al*. 2006)), and *cid-1* (DNA damage checkpoint, whose loss reverts HU-induced developmental arrest and activates *hsp-4p::gfp* (Olsen *et al*. 2006)). Unlike *pri-1* or *pri-2* RNAi, knockdown of none of these genes activated any UPR-ER reporter in somatic cells, except *xpf-1*, whose RNAi weakly induced *Patf-4(uORF)::gfp* at one of two tested timepoints (Supplementary Fig. 5). Notably, in our hands, *cid-1* RNAi failed to induce *hsp-4* (Supplementary Fig. 5), possibly because we initiate RNAi in synchronized L1 stage larvae and not in embryos. In embryos, inactivation of *him-1* activated the UPR-ER (Supplementary Fig. 6), but RNAi of the other tested DNA repair genes did not. We conclude that inactivation of DNA damage response and repair machinery genes doesn’t consistently activate the UPR-ER.

### The UPR-ER is not required to protect the germline against HU-induced replication stress

Because the UPR-ER is activated in somatic cells and embryos after *pri-1* or *pri-2* RNAi, we hypothesized that the UPR-ER protects worms from replication stress and the resulting DNA damage. To test this hypothesis, we used HU, a widely used chemical that inhibits ribonucleotide reductase, which reduces ribonucleosides into deoxyribonucleosides for DNA synthesis (Craig *et al*. 2012). In *C. elegans*, HU exposure leads to S-phase arrest, causing oversized nuclei in the mitotic germline, an extension of the duration of the first cell cycle in early embryos, and germline apoptosis (MacQueen and Villeneuve 2001; Garcia-Muse and Boulton 2005; Stevens *et al*. 2016). To quantify functional requirements of UPR-ER genes in response to replication inhibition, we compared the number of eggs laid per HU-exposed worm during a four-hour recovery period to the number of eggs laid by an unstressed worm of the same genotype, and also after prolonged replication stress by chronic HU exposure from the L1 stage onwards. Acute and prolonged HU exposure both caused fecundity defects, but neither were exacerbated in the tested UPR-ER gene mutants (Fig. 6A-B). Thus, the UPR-ER is apparently dispensable to protect the *C. elegans* germline from replication stress.

**Figure 6.**
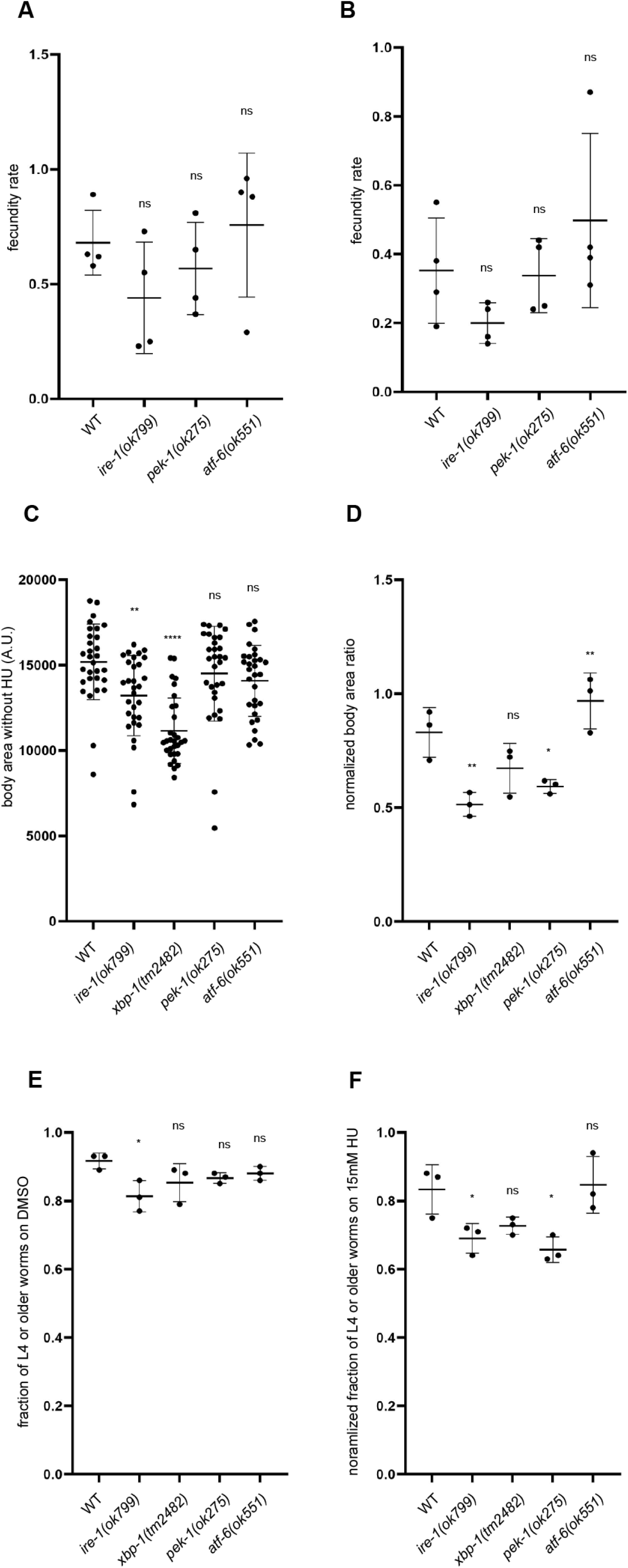
*ire-1* and *pek-1* are required to protect *C. elegans* against HU-induced replication stress. **(A, B)** The graphs show the relative fecundity of wild-type, *ire-1(ok799), pek-1(ok275)*, or *atf-6(ok551)* worms subjected to 20mM HU for 24 hours from late L4 stage (A) or 10mM HU for 72 hours from L1 stage (B). Relative fecundity is calculated as: average number of eggs laid by a HU-treated worm during a 4-hour post-exposure period/average number of eggs laid by an unstressed worm of the same genotype during a 4-hour period. **(C)** Body size quantification of wild-type, *ire-1(ok799), xbp-1(tm2482), pek-1(ok275)*, or *atf-6(ok551)* adult worms grown under unstressed conditions (n=3 experiments totaling >30 animals per genotype). **(D)** The graph shows the normalized average body area ratio of wild-type, *ire-1(ok799), xbp-1(tm2482), pek-1(ok275)*, or *atf-6(ok551)* adult worms, calculated as follows: average body area of worms grown on 15mM HU (n=3 experiments totaling >30 individual HU-treated animals per genotype)/average body area of worms of the same genotype on DMSO. **(E)** The graph shows the fractions of wild-type, *ire-1(ok799), xbp-1(tm2482), pek-1(ok275)*, or *atf-6(ok551)* worms grown past L4 stage on DMSO at 48 hours post-hatching (n=3 experiments totaling >90 individual animals per genotype). **(F)** The graph shows normalized fraction of wild-type, *ire-1(ok799), xbp-1(tm2482), pek-1(ok275)*, or *atf-6(ok551)* worms grown past L4 stage on 15mM HU at 48 hours post-hatching, calculated as follows: fraction of worms past L4 on 15mM HU/fraction of worms past L4 on DMSO (three repeats totaling >120 individual animals per genotype). In all graphs, error bar represents standard deviation. Statistical analysis: ns p>0.05, *p<0.05, **p<0.01, ****p<0.0001 (ordinary one-way ANOVA test corrected for multiple comparisons using Dunnett’s method).

### *ire-1* and *pek-1* are required to protect the soma from HU-induced replication stress

To test whether the UPR-ER protects somatic growth and development from damage caused by prolonged HU exposure, we measured the body area of worms grown on 15mM HU from the L1 stage for 72 hours. Because *ire-1* and *xbp-1* mutant worms have a smaller body size than wild-type worms (Fig. 6C), we normalized body size within each genotype (stressed/unstressed condition). We found that *ire-1* and *pek-1* mutant worms showed a reduced average body size when exposed to HU (Fig. 6D). This suggests that the *ire-1* and *pek-1* branches of the UPR-ER are required for worms to tolerate or resolve prolonged replication stress to achieve normal somatic growth, while the *atf-6* branch is dispensable.

In addition to body size, we also quantified developmental success of worm mutants in the absence of stress and following HU exposure. Loss of *ire-1* caused developmental delay in unstressed conditions, while loss of other UPR-ER components did not (Fig. 6E). When exposed to 15mM HU from the L1 stage on, *ire-1* or *pek-1* mutant worms showed a reduced ability to progress past the L4 stage within 48hr, whereas *atf-6* or *xbp-1* mutation had no effect (Fig. 6F). Collectively, these data show that the *ire-1* and *pek-1* branches, but not the *atf-6* branch, are required to maintain somatic resistance to replication stress.

### Transcriptome analysis of *pri-1* RNAi treated embryos suggests deregulated glycosylation, calcium signaling, and fatty acid desaturation as potential sources of ER stress

To identify genes and processes altered by replication fork stalling, we studied the transcriptomes of wild-type embryos treated with EV or *pri-1* RNAi using RNA-sequencing (RNA-seq). We identified 2785 genes that were up- and 1738 genes that were downregulated following *pri-1* depletion (Fig. 7A; Supplementary Tables 1-3; Supplementary Fig. S7). In line with the above data, *hsp-4* was significantly induced following *pri-1* depletion, as were two other chaperones, *hsp-43* and *hsp-70* (Fig. 7B-D; Supplementary Tables 1-2); others have reported that *hsp-70* is induced by tunicamycin in an *xbp-1* dependent fashion (Urano *et al*. 2002; Lim *et al*. 2014), suggesting that it is an effector chaperone of the UPR-ER. In contrast, neither the mitochondrial UPR chaperones *hsp-6* and *hsp-60*, nor any of the cytoplasmic UPR chaperones of the *hsp-16* family (*hsp-16*.*1, hsp-16*.*11, hsp-16*.*2, hsp-16*.*41, hsp-16*.*48*) were induced (Supplementary Tables 1-3; Fig. 7A). Thus, unbiased transcriptome profiling confirms that the UPR-ER is specifically activated in embryos experiencing replication fork stress, whereas other UPRs are not.

**Figure 7.**
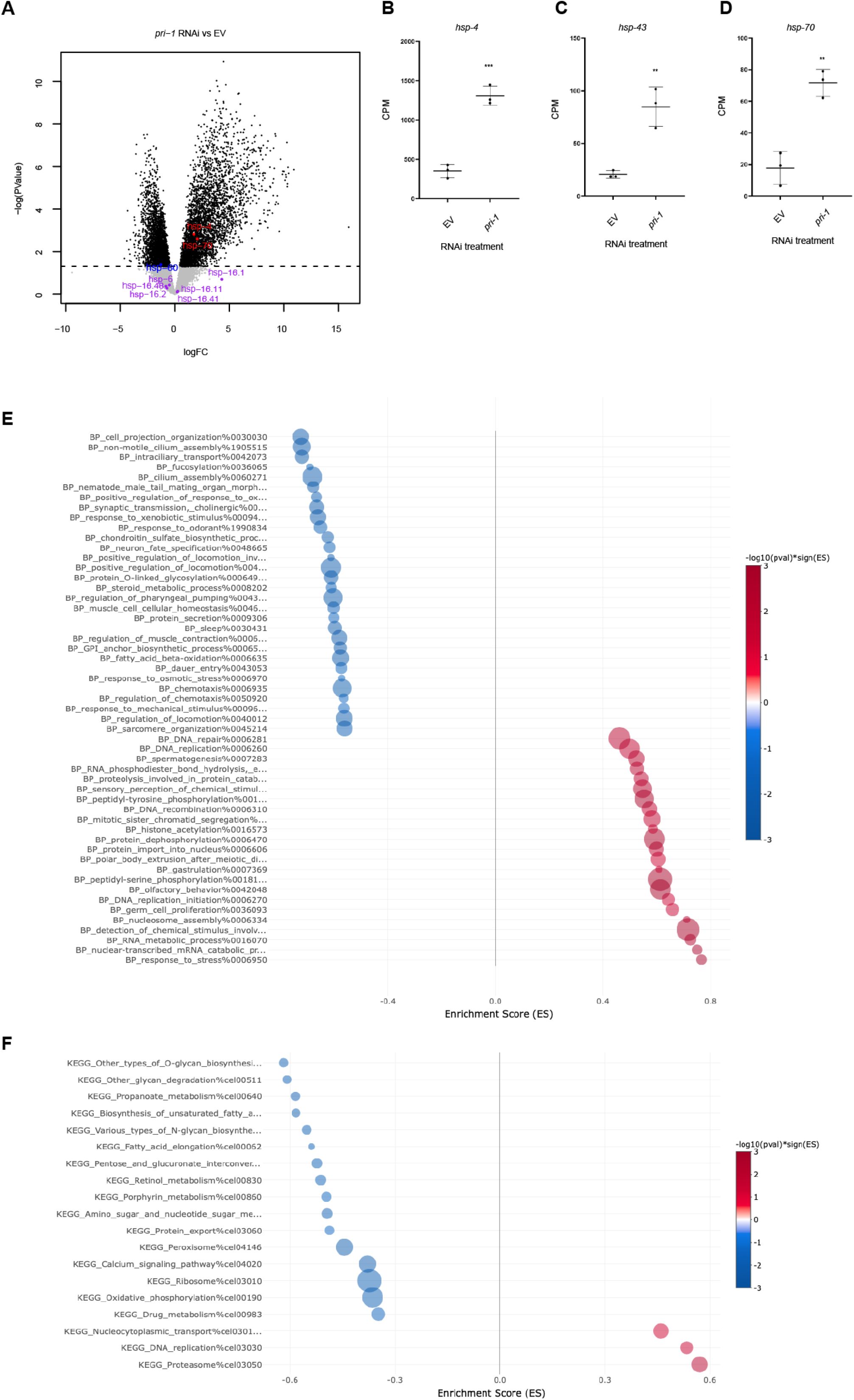
Replication stress in embryos alters protein glycosylation, calcium signaling, and fatty acid desaturation. **(A)** The volcano plot shows the expression of all detected genes in EV and *pri-1* RNAi treated embryos. X-axis, logFC; Y-axis, -log_10_(PValue). Black, PValue <0.05; grey, PValue ≥0.05; blue, highlighted and significantly downregulated; red, highlighted and significantly downregulated; purple, highlighted but not significant. **(B-D)** The graph shows average transcript levels in counts per million (CPM) of *hsp-4, hsp-43*, and *hsp-70* mRNA in EV or *pri-1* RNAi treated embryos; error bars represent standard deviation (n=3 experiments). Statistics: **p<0.01, ***p<0.001 (unpaired Student’s t-test). **(E, F)** The bubble plots show processes enriched negatively (blue) and positively (red) in *pri-1* RNAi treated worms, based on the BP (E) and KEGG (F) databases. Bubbles represent the top 30 or fewer gene sets determined to be statistically significant (cutoffs p<0.05, padj<0.25), as determined by analysis with the easyGSEA function of the eVITTA webserver (Cheng *et al*. 2021). The size the of bubble corresponds to the number of genes represented in each gene set. X-axis: enrichment score (ES).

To delineate how replication fork stress could induce the UPR-ER, we performed Gene Set Enrichment Analysis (GSEA) using the BP and KEGG databases. As expected, terms relating to the UPR-ER stress response were enriched, e.g., the terms “BP_response_to_stress%0006950” (which includes *hsp-4* and *hsp-70*) and “BP_PERK-mediated_unfolded_protein_response%0036499” (Fig. 7E-F, Supplementary Tables S4-S5). Furthermore, we observed an enrichment of terms related to DNA replication and DNA repair (Fig. 7E,F, Supplementary Table S4-5), as expected in worms experiencing replication stress. For example, “KEGG_DNA_replication%cel03030” was one of only three upregulated terms when using the KEGG database for analysis, while upregulated terms identified with the BP database included terms such as “BP_DNA_replication_initiation%000627”, “BP_DNA_repair%0006281”, “BP_DNA_recombination%0006310”, and “BP_double-strand_break_repair_via_homologous_recombination%0000724”. Finally, our analysis identified several processes whose downregulation could indicate the source of UPR-ER activation in *pri-1* RNAi treated worms. Specifically, when using analysis with the KEGG database, of the only 13 downregulated terms, three relate to protein N- and O-glycosylation (“KEGG_Various_types_of_N-glycan_biosynthesis%cel00513”, “KEGG_Other_glycan_degradation%cel00511”, “KEGG_Other_types_of_O-glycan_biosynthesis%cel00514”), one relates to calcium signaling (“KEGG_Calcium_signaling_pathway%cel04020”), and two relate to the biosynthesis of unsaturated fatty acids (“KEGG_Fatty_acid_elongation%cel00062”, “KEGG_Biosynthesis_of_unsaturated_fatty_acids%cel01040”). We conclude that the dysregulation of multiple cellular processes by replication fork stalling after *pri-1* RNAi likely activates the UPR-ER in embryos.

## Discussion

Animals such as *C. elegans* consistently experience and must handle diverse stresses in their environment. Optimal adaptation to such insults requires the deployment of multiple response pathways, and recent studies have identified functional crosstalk between stress responses such as the UPR-ER and the DDR. Here, we show that replication fork stalling strongly induces two branches of the UPR-ER in *C. elegans* embryos. In turn, the UPR-ER is required to protect worms from the deleterious effects of stalled replication forks. Surprisingly, analysis of transcriptional reporters and transcriptome data suggest that it is specifically the UPR-ER that is induced by this stress, whereas other UPRs are apparently not engaged. Our data suggest that replication fork stalling specifically causes ER dysfunction, possibly by disturbing cellular processes that are unique, or especially important, to ER function.

### Replication fork stalling selectively activates the UPR-ER

The UPR-ER, the cytosolic UPR, and the mitochondrial UPR pathways are interconnected adaptive pathways that ensure homeostasis in the face of stress. Conditions that impair general protein folding such as oxidative stress and protein degradation defects induce all three UPRs in *C. elegans* (Rodrigues *et al*. 2011; Hou *et al*. 2014; Bartoszewska and Collawn 2020; Taylor *et al*. 2021). Here, we observed robust induction of two UPR-ER reporters by *pri-1* or *pri-2* RNAi or UV-C irradiation, but did not observe induction of cytosolic or mitochondrial UPR reporters. This lack of induction was confirmed by our unbiased transcriptome profiling. Together, these data indicate that ER proteostasis or membrane lipid equilibrium, but not cytosolic or mitochondrial proteostasis, is disturbed by replication stress. This selectivity appears to rule out a mechanism whereby DNA replication fork stalling causes protein misfolding, aggregation, and proteotoxicity in all organelles. A clue regarding the specific mechanisms underlying UPR-ER activation was revealed by transcriptome profiling following *pri-1* depletion, which revealed dysregulated molecular processes and pathways with important links to the ER. These include protein glycosylation, which is important for the modification of secreted, ER-synthesized proteins; calcium metabolism, which is vital for protein folding in the ER and whose interference via thapsigargin is widely used to study ER protein processing; and fatty acid desaturation, which is essential for maintaining normal ER membrane lipid composition and is monitored by the UPR-ER (Hou *et al*. 2014; Denzel and Antebi 2015; Burkewitz *et al*. 2020; Ho *et al*. 2020; Deng *et al*. 2021; Xu and Taubert 2021). None of these processes are known to impact the activity of the cytoplasmic or mitochondrial UPRs, which may explain why replication fork stalling selectively activates the UPR-ER. However, it remains unclear why genes in these pathways are dysregulated by replication fork stalling.

### Replication stress induces non-canonical, *atf-6*–dependent *hsp-4* expression in embryos

The *hsp-4p::gfp* reporter is widely used in *C. elegans* to monitor activity of the UPR-ER and to infer the presence of ER stress (Calfon *et al*. 2002; Urano *et al*. 2002; Ho *et al*. 2020). Canonical activation of this reporter depends strictly on the IRE-1-XBP-1 pathway. Here, we observed only partially *ire-1*- and *xbp-1*-dependent activation of *hsp-4* after *pri-1* or *pri-2* depletion in embryos. This is surprising because IRE-1 is the only known UPR-ER sensor that processes the unspliced *xbp-1u* mRNA into the mature *xbp-1s* product, which is then translated into XBP-1, the transcription factor that upregulates *hsp-4* expression (Walter and Ron 2011; Gardner *et al*. 2013; Senft and Ronai 2015; Kopp *et al*. 2018; Hetz *et al*. 2020; Xu and Taubert 2021). Because ATF6 is required to express XBP1 mRNA in humans, we studied *atf-6*, which represents the third branch the UPR-ER in *C. elegans* but is thought to be largely dispensable in this organism for stress-induced UPR-ER activity, as many ER stress-activated genes do not require *atf-6* for induction (Shen *et al*. 2005; Lee *et al*. 2007). Interestingly, *atf-6* loss of function significantly diminished *pri-1* or *pri-2* RNAi-induced *hsp-4p::gfp* activation in embryos. In somatic cells, transient *hsp-4* induction that is independent of *ire-1* and *xbp-1* occurs during the differentiation of stem-like seam cells into alae-secreting cells (Zha *et al*. 2019). Although the transcriptional factor B-Lymphocyte–Induced Maturation Protein 1 (*blmp-1*) is required to suppress *hsp-4* in this context (Zha *et al*. 2019), how it is activated is unknown. Our data suggest that *atf-6* may be involved in this process, in line with the view that *atf-6* plays important roles in *C. elegans* development. In sum, our data identify an important new nuance of the mechanisms that fine-tune UPR-ER activation in *C. elegans*.

### *ire-1* and *pek-1* are required for resistance to replication fork stress

A bidirectional crosstalk between the DDR and the UPR-ER has begun to emerge (González-Quiroz *et al*. 2020; Bolland *et al*. 2021). Yeast IRE1 is required for survival on HU (Zha *et al*. 2019). We found that the IRE-1 branch is required to protect *C. elegans* during prolonged HU exposure initiated at an early developmental stage. In contrast, short-term acute HU exposure at a later developmental stage was tolerated, similar to what has been reported with treating *ire-1* worms at L4 stage with *rad-51* RNAi to induce DNA damage (Levi-Ferber *et al*. 2014).

Little evidence exists for roles of the other two branches in response to genotoxicity. We report here that the PEK-1 branch is activated by replication fork stalling and that *pek-1* is required for somatic resistance to HU. In contrast, the *atf-6* branch was not required to protect the soma or germline from HU. This was surprising because, as noted above, *atf-6* is required to induce *xbp-1* in embryos. Nevertheless, our data implicate the UPR-ER as a whole in response to replication fork stalling. Future work will be required to define *ire-1*- and *pek-1*-dependent processes that promote survival and growth in genotoxic conditions.

## Data Availability Statement

The data described in this study are available in the main manuscript, the Supplementary material, or in a public repository. Supplementary Figures S1-S7 are available on GSA Figshare. Supplementary Figures S1, S3, S4, S5 and S6 describe additional experiments using GFP reporters, Supplementary Figure S2 contain additional repeats of Western blots, and Supplementary Figure S7 describes a quality control experiment for the RNA-seq analysis. Supplementary Tables S1-S3 contain lists describing gene expression data identified by RNA-seq; Supplementary Tables S4-5 contain lists describing processes identified by RNA-seq analysis. Supplementary material is available at GENETICS online. Raw and processed RNA-seq files have been deposited at Gene Expression Omnibus accession GSE225569 (https://www.ncbi.nlm.nih.gov/geo/). See methods for information on reagents and strains. *C. elegans* strains described for the first time in this study can be requested from the authors.

## Acknowledgments

We thank Dr. Collin Ewald (Department of Health Sciences and Technology, ETH Zürich, Switzerland) for providing the *Patf-4(uORF)::gfp::unc-54(3’UTR)* strain.

## Supplementary Material

**Figure S1. *hsp-4::gfp* is a faithful reporter for UPR-ER**.

**(A, B)** The figure shows (A) representative micrographs of *hsp-4p::gfp* and *hsp-4::gfp* adult worms, and (B) whole-worm GFP quantification of *hsp-4::gfp* adult worms fed DMSO or 10*µ*g/ml tunicamycin (n=1 experiment, >15 animals per treatment group) for 24 hours. **(C, D)** The figure shows (C) representative micrographs of *hsp-4p::gfp* and *hsp-4::gfp* adult worms, and (D) whole-worm GFP quantification of *hsp-4::gfp* adult worms fed EV or *mdt-15* RNAi (n=2 experiments, >16 individual animals per RNAi group). In all micrographs, the scale bar represents 100*µ*m. In dot plots, each dot represents an individual worm or embryo; error bars represent standard deviation. Statistical analysis: ****p<0.0001 vs. DMSO-treated animals (Welch’s t test).

**Figure S2. *pri-1* or *pri-2* RNAi activates the PEK-1 branch in *C. elegans* embryos**.

Additional repeats (repeats 2 and 3) and full-scale image scan (repeat 1) of immunoblot shown in Fig. 3c. Immunoblots depict the levels of phospho-Ser51 eIF2α (P-eIF2α) and α-tubulin in EV, *pri-1*, or *pri-2* RNAi treated worm embryos (n=3 experiments, for *pri-1(RNAi)* embryos, n=2 experiments for *pri-2(RNAi)* embryos).

**Figure S3. The cytosolic UPR reporter *hsp-16*.*2p::gfp* responds to heat stress in the soma and embryos of *C. elegans***.

**(A, B)** The figure shows (A) representative micrographs of *hsp-16*.*2p::gfp* adult worms, and (B) whole-worm GFP quantification of *hsp-16*.*2p::gfp* adult worms subjected to 1 hour heat shock at 37°C (n=1 experiment totaling 9-10 animals per treatment group). **(C, D)** The figure shows (C) representative micrographs of *hsp-16*.*2p::gfp* embryos, and (D) whole-worm GFP quantification of *hsp-16*.*2p::gfp* embryos subjected to 5 min heat shock at 37°C (n=1 experiment totaling 23-30 embryos per treatment group). In all micrographs, the scale bar represents 100*µ*m. In dot plots, each dot represents an individual animal or embryo; error bars represent standard deviation. Statistical analysis: ****p<0.0001 vs. DMSO-treated animals (Welch’s t test).

**Figure S4. UV-C activates *Patf-4(uORF)::gfp* and *hsp-16*.*2p::gfp* activity in somatic cells.**

**(A, B)** The figure shows (A) representative micrographs, and whole (B) worm GFP quantification of *Patf-4(uORF)::gfp* adult worms irradiated with 400J/m^2^ UV-C (n=3 experiments totaling >20 animals for UV-irradiated and non-irradiated samples). **(C, D)** The figure shows (C) representative micrographs, and (D) whole worm GFP quantification of *hsp-4p::gfp* adult worms irradiated with 400J/m^2^ UV-C (n=3 experiments totaling >20 animals for UV-irradiated and non-irradiated samples). **(E, F)** The figure shows (E) representative micrographs, and (F) whole worm GFP quantification of *hsp-4::gfp* adult worms irradiated with 400J/m^2^ UV-C (n=3 experiments totaling >20 animals for UV-irradiated and non-irradiated samples). **(G, H)** The figure shows (G) representative micrographs, and (H) whole worm GFP quantification of *hsp-6p::gfp* adult worms irradiated with 400J/m^2^ UV-C (n=3 experiments totaling >20 animals for UV-irradiated and non-irradiated samples). **(I, J)** The figure shows (I) representative micrographs, and (J) whole worm GFP quantification of *hsp-16*.*2p::gfp* adult worms irradiated with 400J/m^2^ UV-C (n=3 experiments totaling >20 individual animals for UV-irradiated and non-irradiated samples). In all micrographs, the scale bar represents 100*µ*m. In dot plots, each dot represents an individual animal; error bars represent standard deviation. Statistical analysis: ns p>0.05, *p<0.05, **p<0.001, ****p<0.0001 vs. non-irradiated animals (Welch’s t test).

**Figure S5. RNAi knockdowns of DNA repair pathway genes does not activate the UPR-ER in somatic cells at the late L4 or adult stage**.

**(A-D)** The figure shows (A, C) representative micrographs and (B, D) whole-worm GFP quantification of *hsp-4p::gfp* late L4 worms fed EV, *xpf-1, mus-81, msh-2, him-6, him-1, dog-1*, or *cid-1* RNAi for 48 hours (A, B) or 72 hours (C, D) (n=3 experiments totaling >20 individual animals per RNAi treatment). **(E-H)** The figure shows (E, G) representative micrographs and (F, H) whole-worm GFP quantification of *Patf-4(uORF)::gfp* late L4 worms fed EV, *xpf-1, mus-81, msh-2, him-6, him-1, dog-1*, or *cid-1* RNAi for 48 hours (E, F) or 72 hours (G, H) (n=3 experiments totaling >20 individual animals per RNAi treatment). In all micrographs, the scale bar represents 100*µ*m. In dot plots, each dot represents an individual animal; error bars represent standard deviation. Statistical analysis: ns p>0.05, *p<0.05 vs. EV RNAi-treated animals (Brown-Forsythe and Welch ANOVA test corrected for multiple comparisons using the Dunnett T3 method).

**Figure S6. *him-1* RNAi activates both *ire-1* and *pek-1* branches of the UPR-ER in embryos**

**(A, B)** The figure shows (A) representative micrographs, and (B) GFP quantification of *hsp-4p::gfp* embryos laid by worms fed EV or *him-1* RNAi (n=3 experiments totaling >50 individual embryos per RNAi treatment). **(C, D)** The figure shows (C) representative micrographs and (D) GFP quantification of *Patf-4(uORF)::gfp* embryos laid by worms fed EV or *him-1* RNAi (n=3 experiments totaling >50 individual embryos per RNAi treatment). In all micrographs, the scale bar represents 100*µ*m. In dot plots, each dot represents an individual worm or embryo; error bars represent standard deviation. Statistical analysis: *** p<0.001, **** p<0.0001 vs. EV RNAi-treated animals (Welch’s t test).

**Figure S7. Multidimensional scaling (MDS) plot for EV and *pri-1* RNAi RNA-seq data**.

The figure shows a MDS plot of the distances between gene expression profiles. Distances on the MDS plot correspond to the root-mean-square average of the largest 500 log2-fold-changes between the treatment conditions. Black: EV RNAi; yellow: *pri-1* RNAi.

**Table S1. Expression of genes in EV or *pri-1* RNAi treated embryos as detected by RNA-seq analysis**.

The table lists the gene-level read counts in counts per million (cpm) values for the genes detected in our RNA-seq analysis using the Ensembl transcriptome build WBcel235, including three EV RNAi treated (N2_evi_1, N2_evi_2, N2_evi_3) and three *pri-1* RNAi treated (N2_pri.1i_1, N2_pri.1i_2, N2_pri.1i_3) samples.

**Table S2. Lists of genes significantly upregulated in *pri-1* RNAi treated embryos**.

The table lists all genes that were significantly upregulated in embryos treated with *pri-1* RNAi. Gene: Gene ID; logFC: Log fold change; logCPM: logarithm of counts per million reads; F: value of the F distribution; PValue: p value; FDR: false discovery rate.

**Table S3. Lists of genes significantly downregulated in *pri-1* RNAi treated embryos**.

The table lists all genes that were significantly downregulated in embryos treated with *pri-1* RNAi. Gene: Gene ID; logFC: Log fold change; logCPM: logarithm of counts per million reads; F: value of the F distribution; PValue: p value; FDR: false discovery rate.

**Table S4. Lists of gene sets from the BP database that were up- and downregulated in *pri-1* RNAi treated embryos**.

Details on all gene sets from the BP database that are significantly (pval<0.05, padj<0.25) up- or downregulated in embryos treated with *pri-1* RNAi, as determined by GSEA analysis. Pathway: BP pathway ID; pval: P value; padj; adjusted P-value; ES: enrichment score; NES: normalized enrichment score; size: number of genes in set; genes: names of genes contained in enriched set.

**Table S5. Lists of gene sets from the KEGG database that were up- and downregulated in *pri-1* RNAi treated embryos**.

Details on all gene sets from the KEGG database that are significantly (pval<0.05, padj<0.25) up- or downregulated in embryos treated with *pri-1* RNAi, as determined by GSEA analysis. Pathway: BP pathway ID; pval: P value; padj; adjusted P-value; ES: enrichment score; NES: normalized enrichment score; size: number of genes in set; genes: names of genes contained in enriched set.

